# The extent of B-form structure in a 6S RNA variant that mimics DNA to bind to the *Escherichia coli* RNA Polymerase

**DOI:** 10.1101/242503

**Authors:** Nilesh K. Banavali

**Affiliations:** Laboratory of Computational and Structural Biology Division of Genetics, Wadsworth Center, NYS Department of Health; Department of Biomedical Sciences, School of Public Health State University of New York at Albany

## Abstract

In a recent article by Darst and coworkers, it was found that a non-coding 6S RNA variant regulates a bacterial RNA polymerase by mimicking B-Form DNA, and a few different nucleic acid duplex parameters were analyzed to understand the extent of B-form RNA structure. In this manuscript, a different structural analysis based on conformational distance from canonical A-form and B-form single-strand structures is presented. This analysis addresses the occurrence and extent of both local and global B-form structure in the published 6S RNA variant model.

---

In a recent article describing the structure of non-coding 6S RNA variant bound to an RNA polymerase holoenzyme, Darst and coworkers describe an overall RNA architecture that partially mimics B-form DNA.^1^ In their 3.8 Å single particle cryo-EM structure, they show an average major groove width resembling the B-form, and an average minor groove width resembling the A-form, in the duplex regions of the 6S RNA variant (termed 6S RNA* henceforth) that are long enough to measure groove widths. They also show the pitch and rise parameter values to be nearer those for the B-form, while the sugar pucker, phosphate-phosphate distances, and twist parameter values to be nearer those for the A-form. This analysis of nucleic acid duplex parameters shows that the 6S RNA structure has features of both A-form and B-form architectures.

Proximity to A-form and B-form reference structures can also be assessed using a Root Mean Square Deviation (RMSD) measure,^2^ and we have performed such analysis on known RNA crystal structure conformations of 3 Å or greater resolution at the individual single strand dinucleotide (SSD) level.^3^ This analysis does not require duplex structures, and can assess relative proximity to A- and B-form structures at a local or global level. We showed that RNA SSDs approach a B-form structure in many instances,^3^ and this local B-form structure seems to be achieved through both C2’-endo sugar conformations or base stacking patterns that resemble B-form DNA stacking.^4^ Interestingly, the base RMSD values show a greater propensity for B-form structures than the sugar-phosphate backbone at the SSD level, which suggests an under-representation of B-form-like sugar-phosphate conformations for RNA in the PDB models. While duplex RNA usually adopts an A-form structure with its furanose sugars assuming C3’-endo pucker conformations,^5^ the C2’-endo pucker is present in some functional RNA motifs, and may possibly be implicated in RNA folding.^6,7^

It is possible to apply such RMSD analysis to understand the extent of local and global B-form conformations in the 6S RNA* structure (PDB ID: 5VT0, Chain R). Figure 1A shows the scatter plot of each 6S RNA* SSD conformation in 2D RMSD space for all, base, and backbone non-hydrogen atoms, with the RMSD values computed with respect to canonical A- and B-form SSD reference structures. These RMSD values are normalized by the RMSD difference between the A- and B-form SSD structures, thus setting the RMSD between the A- and B-forms to a value of 1.0, irrespective of the SSD sequence. The interpolation line between the A- and B-forms is shown as a dotted blue line. The dividing solid red line separates the conformational space into two regions, one nearer to the B-form (bordered by a dotted cyan line), and another closer to the A-form (bordered by a dotted pink line). If all SSDs are simply separated into ones relatively closer to the A-form or the B-form, then a majority of the SSDs are clustered near the canonical A-form, but some SSDs clearly cross over to the B-form, mostly in conformational regions further away from the canonical B-form. This behavior was also observed in our larger crystallographic survey,^3^ such that crossing over to the B-form region seems to be easier for RNA if it moves away from both canonical forms, and the region very close to the canonical B-form structure does not seem to be accessible.

**Figure 1:**
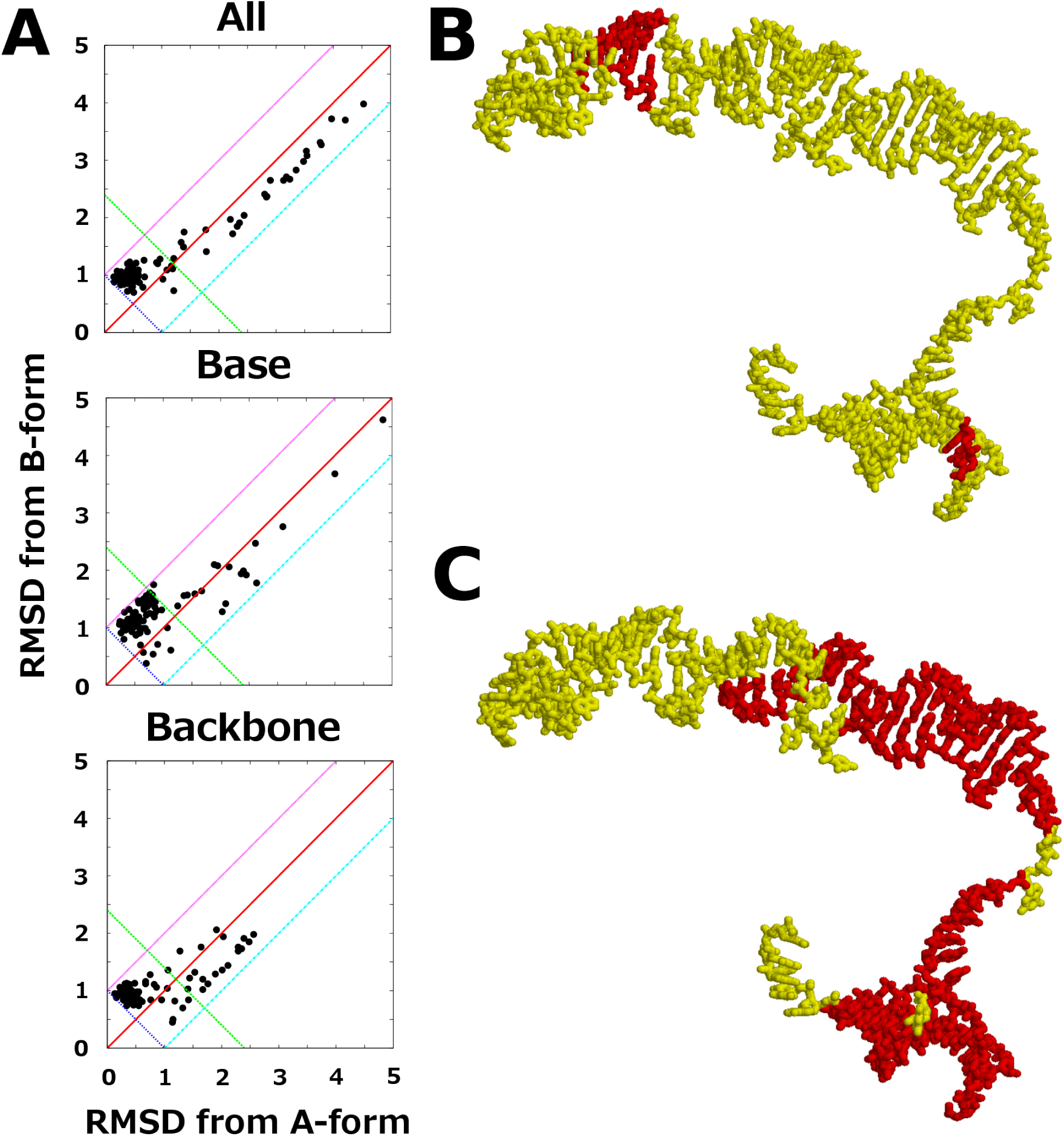
Analysis of SSD and longer RNA single strand B-form conformations in the 6S RNA variant structure bound to the *E. coli* RNA polymerase. (A) SSD conformations scatter plots for all SSD non-hydrogen atoms (top), base SSD non-hydrogen atoms (center), and sugar-phosphate backbone SSD non-hydrogen atoms (bottom) in normalized 2D RMSD space from the canonical A-form and B-form structures. The interpolation line between the two canonical forms is in dotted blue, the dividing line between A- and B-form regions is in solid red, the B-form region border is in dotted cyan, and the A-form region border is in dotted pink. (B) Structural location of SSDs (red) near the canonical B-form structure, as judged by presence in the B-form half and an additional cutoff of 1.0 in the perpendicular distance from the dotted blue interpolation line. (C) Indication of overall B-form segments (red) of extended stacked single strand regions of the 6S RNA structure according to base non-hydrogen atom RMSD values.

To see which SSDs of the 6S RNA* structure might assume a B-form conformation reasonably close to the canonical B-form, a normalized RMSD cutoff was used to limit the distance from the interpolation line (1.0 in this case, shown as a green dotted line). The conformations with base non-hydrogen atom RMSD values that fall in the B-form region within this cutoff distance are shown in red in Figure 1B. This analysis shows that, even for the base moieties more likely to show B-form conformations, there are only six B-form SSD conformations distributed throughout the entire RNA structure that are nearer the canonical B-form. To assess the A-form or B-form proximity for longer stretches of the RNA strands, the single stranded segments that had continuous stacking were identified. These stretches consisted of residues 32–42, 43–47, 58–68, 72–81, 83–85,99–107, 112–134, 137–142, and 144–153. The RMSD values for different components of these single stranded regions are listed in Table 1. The stacked single strand segments showing greater proximity to the B-form than the A-form at these longer length scales (for the base non-hydrogen atoms) are shown in red in Figure 1C. It can be seen that a substantial part of the 6S RNA variant structure assumes overall single stranded conformations that resemble the B-form more than the A-form.

**Table 1:** Canonical form proximity of stacked single strand segments of the 6S RNA variant structure.

| Residues | $D_{RMSD}$<br>All | $D_{RMSD}$<br>Base | $D_{RMSD}$<br>Backbone | $RMSD_A$<br>All | $RMSD_B$<br>All | $RMSD_A$<br>Base | $RMSD_B$<br>Base | $RMSD_A$<br>Backbone | $RMSD_B$<br>Backbone |
| --- | --- | --- | --- | --- | --- | --- | --- | --- | --- |
| 32-42 | -0.28 | 0.12 | -0.40 | 0.50 | 0.78 | 0.70 | 0.59 | 0.42 | 0.82 |
| 43-47 | -0.60 | -0.61 | -0.60 | 0.45 | 1.05 | 0.60 | 1.22 | 0.43 | 1.03 |
| 58-68 | -0.15 | 0.10 | -0.23 | 0.74 | 0.89 | 0.89 | 0.79 | 0.69 | 0.92 |
| 72-81 | -0.34 | -0.21 | -0.39 | 0.51 | 0.85 | 0.73 | 0.95 | 0.44 | 0.83 |
| 83-85 | -0.41 | -0.14 | -0.46 | 1.19 | 1.60 | 2.00 | 2.15 | 1.08 | 1.54 |
| 99-107 | -0.48 | -0.31 | -0.52 | 0.45 | 0.93 | 0.56 | 0.87 | 0.42 | 0.94 |
| 112-134 | -0.14 | 0.09 | -0.25 | 0.54 | 0.68 | 0.64 | 0.55 | 0.49 | 0.74 |
| 137-142 | -0.34 | 0.07 | -0.42 | 0.64 | 0.98 | 1.27 | 1.20 | 0.53 | 0.95 |
| 144-153 | -0.27 | 0.04 | -0.35 | 0.55 | 0.83 | 0.70 | 0.67 | 0.51 | 0.86 |

$D_{RMSD} = RMSD_A - RMSD_B$ , where $RMSD_A$ and $RMSD_B$ are RMSDs of a particular structure from the canonical A- and B-forms, respectively. Structures in the A-form half of 2D RMSD space have negative $D_{RMSD}$ values, and structures in the B-form half have positive $D_{RMSD}$ values. All RMSD values are calculated after a Kabsch least squared fit<sup>8</sup> of the two structures using either all non-hydrogen atoms (All), or using only the non-hydrogen atoms of its base moieties (Base), or using only the non-hydrogen atoms of its sugar-phosphate backbone (Backbone). All RMSD values are normalized by the corresponding difference in RMSD between the canonical A- and B-form reference structures.

The proportion of the 6S RNA* SSDs merely in the B-form half of the 2D RMSD space is less than 35% of the structure, while that closer to the canonical B-form is about 5% of the structure. In contrast, the proportion of SSDs merely in the A-form half (68%) is similar to the proportion close to the canonical A-form (60%). This suggests that the 6S RNA* can mimic DNA by having only a third of its SSDs resembling the B-form more than the A-form, and only a twentieth of its SSDs being nearer the canonical B-form structure. The propensity of B-form structure in RNA crystal structure SSD conformations that we assessed previously using the base non-hydrogen RMSD is also near 5%.^3^ This 6S RNA* structure therefore suggests that RNA may be able to mimic DNA functionally by crossing over into the B-form 2D RMSD space, even through just a substantial minority of its SSDs assuming conformations closer to the B-form than the A-form.

## References

[1] Chen, J.; Wassarman, K. M.; Feng, S.; Leon, K.; Feklistov, A.; Winkelman, J. T.; Li, Z.; Walz, T.; Campbell, E. A.; Darst, S. A. 6S RNA Mimics B-Form DNA to Regulate Escherichia coli RNA Polymerase. Molecular Cell 2017, 68, 388–397.

[2] Banavali, N. K.; Roux, B. Free energy landscape of A-DNA to B-DNA conversion in aqueous solution. Journal of the American Chemical Society 2005, 127, 6866–6876.

[3] Sedova, A.; Banavali, N. K. RNA approaches the B-form in stacked single strand dinucleotide contexts. Biopolymers 2016, 105, 65–82.

[4] Sedova, A.; Banavali, N. K. Geometric Patterns for Neighboring Bases Near the Stacked State in Nucleic Acid Strands. Biochemistry 2017, 56, 1426–1443.

[5] Fuller, W.; Hutchinson, F.; Spencer, M.; Wilkins, M. Molecular and crystal structures of double-helical RNA: I. An X-ray diffraction study of fragmented yeast RNA and a preliminary double-helical RNA model. Journal of Molecular Biology 1967, 27, 507–524.

[6] Gherghe, C. M.; Mortimer, S. A.; Krahn, J. M.; Thompson, N. L.; Weeks, K. M. Slow conformational dynamics at C2´-endo nucleotides in RNA. Journal of the American Chemical Society 2008, 130, 8884–8885.

[7] Mortimer, S. A.; Weeks, K. M. C2´-endo nucleotides as molecular timers suggested by the folding of an RNA domain. Proceedings of the National Academy of Sciences 2009, 106, 15622–15627.

[8] Kabsch, W. A discussion of the solution for the best rotation to relate two sets of vectors. Acta Crystallographica Section A: Crystal Physics, Diffraction, Theoretical and General Crystallography 1978, 34, 827–828.

